# Germline DNA replication timing shapes mammalian genome composition

**DOI:** 10.1101/258426

**Authors:** Yishai Yehuda, Britny Blumenfeld, Nina Mayorek, Kirill Makedonski, Oriya Vardi, Yousef Mansour, Hagit Masika, Marganit Farago, Shulamit Baror-Sebban, Yosef Buganim, Amnon Koren, Itamar Simon

## Abstract

Mammalian DNA is replicated in a highly organized and regulated manner. Large, Mb-sized regions are replicated at defined times along S phase. DNA Replication Timing (RT) has been suggested to play an important role in shaping the mammalian genome by affecting mutation rates. Previous analyses relied on somatic DNA RT profiles, while to fully understand the influences of RT on the mammalian genome, germ cell RT information is necessary, as only germline mutations are passed to offspring and thus affect genomic composition. Using an improved RT mapping technique that allows mapping the RT from limited amounts of cells, we measured RT from two stages in the mouse germline - primordial germ cells (PGCs) and spermatogonial stem cells (SSCs). The germ cell RT profiles were distinct from those of both somatic and embryonic tissues. The correlations between RT and both mutation rate and recombination hotspots were not only confirmed in the germline tissues, but were shown to be stronger compared to correlations with RT of somatic tissues, emphasizing the importance of using RT profiles from the correct tissue of origin. Expanding the analysis to additional genetic features such as GC content, transposable elements (SINEs and LINEs) and gene density, also revealed a stronger correlation with the germ cell RT maps. GC content stratification along with multiple regression analysis revealed the independent contribution of RT to SINE, gene, mutation and recombination hotspot densities. Taken together, our results point to the centrality of RT in shaping multiple levels of mammalian genome composition.

## Introduction

DNA replication follows a highly regulated temporal program consisting of reproducible RT of different genomic regions (Goren and Cedar 2003; MacAlpine et al. 2004; Norio et al. 2005; Schwaiger and Schubeler 2006; Karnani et al. 2007; Farkash-Amar et al. 2008; Desprat et al. 2009; Hiratani et al. 2009; Schwaiger et al. 2009). RT is conserved across species (Farkash-Amar et al. 2008; Ryba et al. 2010; Yaffe et al. 2010; Pope et al. 2012), and within a species about 50% of genomic regions have stable RT across cell types, while the other 50% have variable RT between cell types (Hiratani et al. 2010; Rivera-Mulia and Gilbert 2016). The importance and role of this temporal organization are still unclear.

RT correlates with many genomic and epigenomic features including transcription (Braunstein et al. 1982; Gilbert 1986; Farkash-Amar et al. 2008; Hiratani et al. 2008), gene density (Cohen et al. 1998), chromatin state (Farkash-Amar and Simon 2010; Farkash-Amar et al. 2012), retrotransposon density (Woodfine et al. 2004; Hiratani et al. 2008), lamina proximity (Farkash-Amar et al. 2012), topological state (Pope et al. 2014; Dileep et al. 2015; Kenigsberg et al. 2016), and GC content (White et al. 2004; Woodfine et al. 2005; Farkash-Amar et al. 2008; Kenigsberg et al. 2016). RT is also associated with mutation rates both in cancer (Donley and Thayer 2013; Lawrence et al. 2013) and in the germline (Stamatoyannopoulos et al. 2009; Chen et al. 2010). Late replicating regions are enriched with point mutations (Chen et al. 2010; Cui et al. 2012), whereas the association between CNVs and RT is more subtle and depends on the mechanism of CNV generation (Koren et al. 2012) and on the organism (reviewed in (Blumenfeld et al. 2017)). We recently investigated the correlation between RT and GC content and found that different substitution types have different associations with RT: late-replicating regions tend to gain both As and Ts along evolution. whereas early replicating regions tend to lose them (Kenigsberg et al. 2016). Measuring the levels of free dNTPs at different time points along S phase revealed an increase in the dATP+dTTP to dCTP+dGTP ratio along S, suggesting that a replication timing-dependent deoxynucleotide imbalance may underlie this mutation bias.

The association between RT and germline mutation frequency points to the importance of RT in shaping the genome sequence. To fully understand this association would require profiles of replication timing in germ cells. However, all previous studies used somatic tissue RT profiles as proxies for the investigation of the evolutionary impacts of RT. Thus, it is crucial to measure the RT in germ cells.

Germ cells refer to all the cells in an organism that pass on their genetic material to progeny. Mouse oogenesis and spermatogenesis involve 25 and 37- 62 cell divisions, respectively (Drost and Lee 1995). Mutations occurring at each step of this process are inherited by the next generation and thus all steps in this process are important from an evolutionary standpoint. RT has been measured in the early stages of this process (ESC to EpiSC (Hiratani et al. 2010)), but there is no data regarding replication timing at later stages during which the majority of cell divisions occur (Drost and Lee 1995) and during which a high percentage of germline mutations likely accumulate. In order to start filling this gap, we have measured RT at two different stages along the germline: primordial germ cells (PGCs, isolated directly from gonads of E13.5 mouse embryos) and spermatogonial stem cells (SSCs, isolated directly from testes of p5 pups). The main limitation for measuring RT in these stages is the small amount of available cells. The current methods for measuring genome wide RT (reviewed in (Gilbert 2010) and (Farkash-Amar and Simon 2010)), are usually applied on millions of growing cells (Farkash-Amar et al. 2008; Dileep et al. 2012), which is not feasible for many cell types including in vivo germ cells.

By improving the RT mapping method, we were able to generate reliable RT maps from as few as 1000 S-phase cells. We first demonstrated the reliability of this method on small populations of mouse embryonic fibroblasts (MEFs). We then measured the RT of in vivo PGCs and of isolated SSCs. RT patterns of germ cells were highly correlated to each other, and were more similar to early embryonic tissues than to somatic cells. Both germline mutation and recombination hotspot densities correlated more strongly with the RT of the germ cell compared to that of somatic tissues, as expected. Mapping RT in the germline enabled us to similarly explore other genomic features such as GC content, LINEs, SINEs and gene density, all of which correlated better with germ cell RT. GC content stratification, as well as multiple regression analyses revealed that germ cell RT contributes to SINE, gene and recombination hot spot densities as well as to mutation rates, independently from the contributions of GC content. Taken together, our results suggest a role for germ cell RT in shaping multiple features of the genome sequence.

## Results

### RT maps from small amounts of celsl

RT maps are usually generated from at least 100,000 S-phase cells (Yehuda et al. 2017). In order to measure RT in germ cells, we first established our ability to map RT from as few as 1000 S-phase cells. To this end, we optimized the RT profiling technique to minimize cell loss by optimizing fixation conditions, avoiding material transfer between tubes, using a slow flow rate during cell sorting and optimizing DNA extraction and library preparation protocols (see supplementary methods). Using this improved technique, we measured RT of MEFs using 10^3^, 10^4^ or 10^5^ S-phase cells. Triplicates were highly similar for 10—4 and 10—5 samples (R>0.91 and R>0.95, respectively) and quite similar even in the 10—3 triplicates (R>0.76). Moreover, the RT maps from all cell numbers were quite similar to each other and to published RT map (Hiratani et al., 2010) and distinct from RT profiles generated from other tissues (**Figure 1)**.

**Figure 1.**
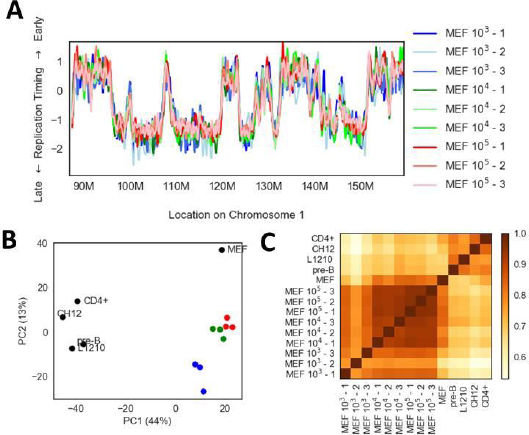
RT mapping of small populations of cells. **A)** RT maps of 10^3^,10^4^ and 10^5^ MEF cells, in triplicates, along a ~80 Mb region on chromosome 1. **B)** PCA analysis of RT profiles. Plot of PC1 vs PC2 for RT profiles of multiple MEF samples (described in A or published) and other somatic cells either sequenced by us (L1210 and PreB) or published (CH12 and CD4). The MEF samples mapped in this paper are color coded as in A**. C)** A heatmap of spearman correlation coefficients between different RT profiles.

Moreover, autocorrelation analyses performed on the different samples were almost identical (**Figure S1**). Taken together, our results demonstrate the ability to obtain reliable RT maps from as little as 1000 S phase cells.

## Replication timing profiles of the mammalian germline

To evaluate germ cell RT we concentrated on the two stages in mouse germline development in which most germ-cell divisions occur: PGC and SSC (**Figure 2A**). We isolated 1,000 to 10,000 G1 and S phase cells in triplicates from PGCs (both male and female) and SSCs, and generated RT maps (see methods). Despite the small amounts of cells used, PGC and SSC RT maps showed high reproducibility (R>0.8 and R>0.85, respectively) and correlated with many genomic features (**Figure S2**), as had been shown for other RT maps (Farkash-Amar and Simon 2010). Interestingly, we found high similarity between the different PGC samples regardless of their gender (**Figure S3**). Moreover, as expected, PGC and SSC RT showed specific association with chromatin accessibility in germ cells (**Figure S4**), further supporting their accuracy.

**Figure 2.**
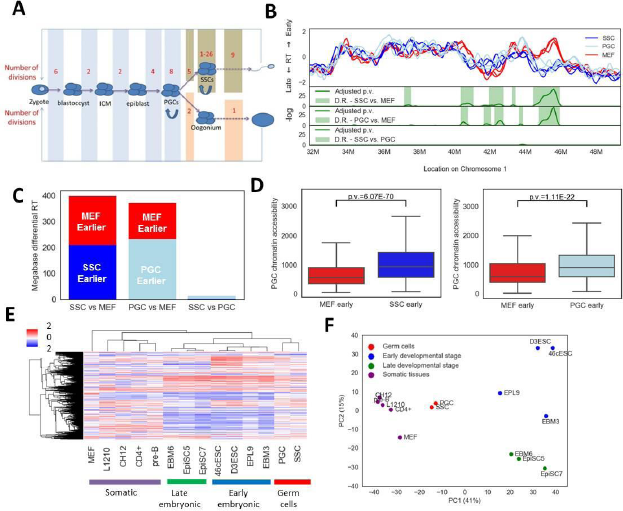
Germ cell RT. **A)** Schema of the germline including the main stages from the zygote to the gonads, both for male and female mice. The number of cell divisions in each stage is shown (data taken from (Drost and Lee 1995)). **B)** RT map of triplicates of PGCs, SSCs and MEFs (10^5) along ~16Mb region of mouse Chr1. Below, three graphs are shown depicting the Bonferroni-corrected p values of the likelihood ratio test for a pairwise comparison between two cell types. Differential regions are highlighted in green. **C)** Bar graphs showing the total size (in megabases) of the differential regions, each bar graph is divided into two portions depicting the size of the regions that are earlier in the SSC (Blue), PGC (light blue) and MEF (red). **D)** Boxplots showing the distribution of PGC chromatin accessibility data in regions showing differential RT between SSC or PGC and MEF. Chromatin accessibility distribution was separated into MEF early versus germ cells early. P values (two sided t test) are shown above the box plots. **E)** RT profiles from the current work along with published embryonic tissues (Hiratani et al. 2010) were hierarchically clustered. Only switching RT regions (see methods) were included. **F)** PCA of RT profiles, plot of PC1 vs PC2 for RT profiles of different types, using the same color code and regions as in E.

The PGC and SSC profiles were very similar to each other (R=0.86), but showed significantly less similarity to the MEF RT profile (R= 0.74 for both PGC and SSC) (**Figures 2B-C and S2B**). We identified statistically significant differential RT regions (see methods) between SSCs, PGCs and MEFs (**Figure 2B-C**). Overall, we found approximately 400Mb and 370Mb of differential RT between MEFs and SSC or PGC, respectively (**Figure 2C**). On the other hand, the SSC and the PGC profiles were very similar (**Figure 2B-C**), with only 14 Mb of differential RT. We confirmed the accuracy of these differential regions by analyzing their chromatin accessibility using published PGC data (Guo et al. 2017). Indeed, early-replicating regions in PGC or SSC were significantly more accessible than regions replicating later in the germ cells (**Figure 2D**).

In order to put the germ cell RT maps in a broader context, PGCs and SSCs were compared to many published RT maps, expanding the work of Hiratani et al. (Hiratani et al. 2010). As was previously reported, embryonic tissues RT clustered into early and late embryonic stages (Hiratani et al. 2010). The germ cells clustered as a third embryonic cells cluster, distinct from terminally differentiated cells (**Figures 2E and 2F**).

### Mutation rate and recombination hotspot density correlate most strongly with the RT of germ cells

Although the mechanism(s) responsible for the association between mutation rates and replication timing is still under investigation, it is clear that it stems from differences between early and late replicating regions, either in mutation rates directly or in DNA repair rates (Blumenfeld et al. 2017). As inter-mammalian divergence reflects germline mutation rates, we expected to find a stronger correlation with germ cell RT than with somatic cell RT. Indeed, we found stronger correlations between inter-mammalian divergence and PGC or SSC RT than MEF RT (R=-0.63 and E0.65 versus E0.52; **Figure S5A**). To further emphasize this trend, we used published data that divided the mouse genome into two types of regions – those that show similar RT across 28 mouse RT datasets (constitutive RT) and those that show variability between tissues (developmental or switching RT) (Hiratani et al. 2010). As expected, in switching RT regions the correlation was much stronger in germ cells than in somatic cells (**Figure 3A-C**). Further analysis of the differential regions between MEFs and germ cells revealed that Germ-Early MEF-Late regions have significantly lower mutation rates than Germ-Late MEF-Early regions (**Figures 3D**), further demonstrating that the mutation rates follow germ cell RT more strongly than somatic cell RT.

**Figure 3.**
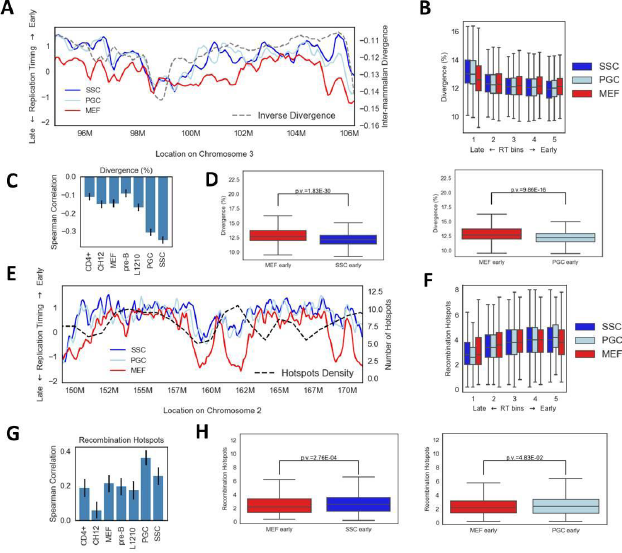
Mutation rate and recombination hotspots correlate better with germ cell RT. The stronger association between inter-mammalian divergence (A-D) and recombination hot spots (E-H) with germ cell RT is shown as RT maps (A, E), box plots in 5 RT bins (B, F), and bar graphs capturing the Spearman correlation coefficients along with confidence intervals (bars) for multiple cell types (C,G) and boxplots (as in Figure 2D) showing the distribution in differential regions (D,H). B, C, E, and F were calculated using only the switching RT regions of the genome (Hiratani et al. 2010).

Another germ cell related feature is meiotic recombination hotspots (Lange et al. 2016). In order to analyze its association with germ cell RT, we took advantage of the recently published dataset depicting the genome-wide recombination hotspots using Spo11 pull-down in mouse sperm cells (Lange et al. 2016). Analyzing the recombination hot spots data (in 1Mb windows) revealed a stronger correlation with germ cell RT for both PGC and SSC (**Figures 3E-H and S5B**) compared to somatic cell RT. Taken together, these results emphasize the centrality of replication timing in determining germline mutation and recombination rates, and establish a resource for further studies of the influence of replication timing on germline genetic and epigenetic events.

### Germline RT is associated with GC content and gene and transposon densities

Having demonstrated that germline RT provides the best proxy, so far, for germline mutation and recombination rates, we turned to search for additional genetic properties that specifically relate to germline RT. Finding such features, would suggest that they originated in the germ cells probably as a consequence of RT. On the other hand, finding a feature that is associated with the RT in all tissues to the same extent, would suggest that this feature is most probably affecting the RT and thus it has the same effect in all tissues.

We explored four additional genomic features that are known to be associated with RT but for which the causative relationship with RT has been unclear – GC content, and SINE, LINE and gene density. Early replicating regions tend to have higher GC content (Woodfine et al. 2004; Farkash-Amar et al. 2008). LINEs are known to populate mainly late replicating regions (Hiratani et al. 2008), whereas SINEs and genes are known to populate mainly early replicating regions (Woodfine et al. 2004; Woodfine et al. 2005). We have previously shown that the genomic GC distribution (GC content) depends on RT, in a mechanism by which RT affects the type of mutations that occur at early and late S (Kenigsberg et al. 2016). According to this explanation, we expect to obtain higher correlations to GC content when using germ cell RT data. Indeed, using the same strategy as with mutation rates, we found stronger correlations of GC content with germ cell RT than with somatic cells RT (**Figures 4A and S6**). Interestingly, using the same approach, we found that LINE, SINE and gene density also correlate better with RT in germ cells (**Figures 4B-D** and **S6**). Thus, our findings suggest that it is less likely that either gene or retrotransposon densities affect RT, but rather point to an influence of RT on those features through a germline-related mechanism.

**Figure 4.**
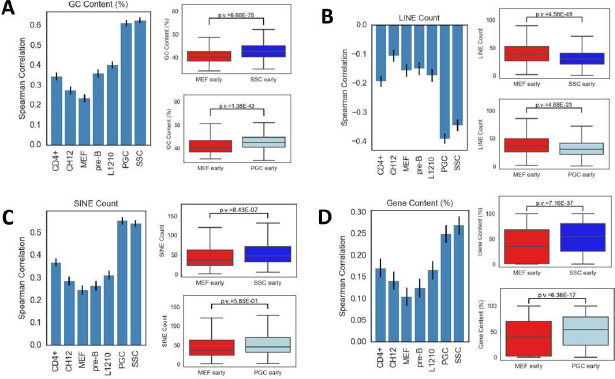
Cell-type specific association of RT with additional genomic features. RT association with GC content (A), LINE counts **(B)**, SINE counts **(C)** and gene coverage **(D)** in 100kb windows. **Left:** bar plots showing the spearman correlation coefficients (along with confidence intervals) with the RT of different cell types, in the switching RT regions of the genome; **Right**: box plots (as in Figure 2D) showing the distribution of these features in differential RT regions.

### RT directly associates with SINE, gene, mutation and recombination hotspot densities

Analyzed the association between RT and each genomic feature in each bin (**Figure 5A)**. We found that for LINE density, the contribution of RT was small relative to the contribution of GC content. On the other hand, for SINE density, mutations rates and recombination hotspots density, germ cell RT was a major contributor even after accounting for GC content. Gene density showed an intermediate pattern in which RT contributed only in genomic regions of low GC content, and was not important in other parts of the genome.

To corroborate this point further, we built a multiple regression model, which allowed us to see the additional contribution of RT over the contribution of GC content. We simultaneously built two models either starting with GC content or with germ cell RT. These models revealed that for LINE the additional contribution to the percent variance explained (PVE) of RT beyond GC content was very small. On the contrary, when predicting SINE density, gene density, mutation rate and recombination hotspots density, adding RT as a predictor increased the PVE by a factor of 20%, 25%, 34% and 35%, respectively, relative to the PVE from using only GC content (**Figure 5B**). Further confirmation of this conclusion was obtained by partial correlation analysis (Figure S8). Taken together, these results demonstrated the independent association between RT and multiple genomic features, suggesting it may has a causative role in their formation.

**Figure 5.**
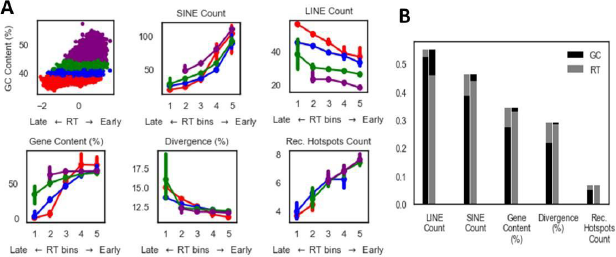
Independent association of RT with genomic features. **(A)** Scatter plot showing the association between RT and GC content and its stratification into four GC content groups; associations between multiple genomic features and RT stratified by GC content, are shown using the same colors as in the scatter plot. **(B)** Barplots depicting relative contribution of RT and GC content to percent variance explained for LINE, SINE, gene coverage, mutation rate and recombination hot spots. For each predicted feature, the model was created twice beginning with either RT or GC content. The order of the addition of the predictors to the model is from bottom to top. Similar results were obtained with SSC RT data (Figure S7).

## Discussion

By improving the RT profiling technique, we were able for the first time, to map the RT of two stages of the mouse germline. We have profiled both E13.5 PGCs and *invitro*-grown SSCs, and found that their RT profiles are similar. Our results add a new dimension to previous efforts to map the RT of multiple mouse developmental stages (Hiratani et al. 2010). We found that the two stages of germ cell development clustered together, and to a lesser extent, clustered with other embryonic tissues while remaining distinct from terminally differentiated cells. The similarity between the RT maps of PGCs and embryonic tissues is not surprising since PGCs are taken from an early embryonic stage prior to terminal differentiation. Though SSCs are isolated from young mice and therefore may reflect later developmental stages, our finding of their similarity to embryonic stages may reflect their stemness (Komeya and Ogawa 2015) resembling many of the embryonic stages examined. Further research is needed to expand our study to additional germ cells models (Geijsen et al. 2004; Seandel et al. 2007; Hikabe et al. 2016; Mitsunaga et al. 2017), which reflect other stages in germ cells development.

It is well established now that RT is associated with both germline and somatic mutations (reviewed in (Blumenfeld et al. 2017)), however, due to a lack of information regarding germ cell RT, previous studies of germline mutations used somatic cell replication profiles as a proxy. By profiling RT in germ cells, we showed that the correlation between mutation rate and RT is stronger than when using somatic cell RT profiles. More generally, obtaining germline-specific RT data is very important for understanding the regional variation in mutation rate (RViMR) along the genome (Hardison et al. 2003; Hodgkinson and Eyre-Walker 2011; Makova and Hardison 2015). It has been shown that RViMR is dependent mainly on RT and transcription activity (Lawrence et al. 2013), which both differ between tissues. Using the correct tissue data improves RViMR estimation (Polak et al. 2015; Supek and Lehner 2015) and accordingly, germ cell RT data is especially important for estimation of germline RViMR. Obtaining a correct cancer related RViMR turned out to be crucial for the identification of new cancer-associated genes (Lawrence et al. 2013). Similarly, obtaining a reliable germline RViMR is important for understanding the selection forces acting on various genes (Caporale 2000; Martincorena and Luscombe 2013), for interpreting the importance of genetic variation and *de novo* mutations for diseases (Samocha et al. 2014), and for reliably performing inter species alignments (Li and Miller 2003). Performing similar experiments in human germ cells will be even more informative, since i) there is more data regarding mutations and CNVs in humans than in mice, and ii) getting a better estimate for the local mutation frequency in humans may allow better understanding of disease-related mutations.

We found that the strongest correlations for germline mutation rate and for recombination hotspots density are found with germ cell RT profiles. This suggests that the strength of the correlation is indicative of the tissue of origin of the studied association. Indeed, correlation between RT and an epigenetic feature (like chromatin accessibility) is found to be stronger when both the RT and the chromatin accessibility data are from the same tissue (**Figure S4**).

We found the strongest correlation between RT and GC content in germ cells, supporting our previous finding that the mutation spectrum in genomic late replication domains shapes mammalian GC content (Kenigsberg et al. 2016). Expanding this idea to other genomic features such as SINE, LINE, and gene density, revealed that all those features correlate more strongly with germ cell RT profiles, suggesting that germline tissues are indeed the relevant tissues of origin for these correlations. This finding implies that it is less likely that these features are involved in affecting RT, either directly or indirectly, since in that case we would expect them to influence RT in all tissues similarly. Rather, our findings suggest that the association of these features with RT occurs in the germline. Nevertheless, it does not necessarily imply that RT is directly affecting these features, since it can be that other processes, associated with RT, like certain chromatin modifications, chromatin accessibility, or the association of certain proteins with chromatin in germ cells, are the direct effectors. Reliable chromatin data from germ cells is required for further evaluating this point.

Previous work has shown that the strongest correlation between RT and both GC content and retroelement density is obtained when using ectoderm tissue RT profiles (Hiratani et al. 2010), but the reason for this phenomenon remained obscure. Our results explain this finding, since germ cells are from ectoderm origin and thus their RT maps are similar (**Figure S9**).

By using the germ cell RT data we were able, for the first time, to address the relative contribution of RT and GC content to multiple genomic features. Interestingly, we found an independent contribution of RT to all examined genomic features besides LINE density. It was shown that L1 elements (LINE) are associated with AT-rich, late-replicating regions, whereas Alu elements (SINE) are associated with GC rich, early-replicating regions. Detailed analysis of old and new SINE and LINE elements revealed that both integrate preferably into AT rich regions, but SINEs are preferentially deleted from those regions and thus old SINEs are enriched in high GC, early replicating regions whereas new SINEs are enriched in low GC, late replicating regions (Lander et al. 2001; Deininger and Batzer 2002). Our results, showing GC-content independent RT association only with SINEs but not LINEs densities, suggest that RT plays a role in the deletion process rather than in the integration process. This conclusion is consistent with the finding that both point mutations and deletions are more prevalent in late replicating regions (Blumenfeld et al. 2017).

The independent association between mutation rate and RT has been reported before using somatic cells RT data (Chen et al. 2010). Our new germ cell RT data confirms previous results and further demonstrates the importance of RT in determining germline mutation rates.

The association between recombination hot spots and somatic cells RT was studied in humans and revealed a stronger association in females than in males (Koren et al. 2012). This study estimated recombination events by analyzing 15,000 Icelandic parent-offspring pairs. Using a direct measurement of the locations of the DNA recombination associated double strand breaks (Spo11 oligo mapping) in mouse sperm cells, we were able to show a strong correlation between germ cell RT and recombination hotspots density (in 1Mb windows). Interestingly, our results differ from the previous report in two aspects – i) we found a much stronger correlation with male recombination hotspots than previously reported. ii) In the current study, we found that RT contributes to recombination hotspots even after controlling for GC content whereas the previous report suggested that the association between RT and recombination strongly depends on GC content. Differences between the studies in i) the organism studied (mouse versus human); ii) the source of the RT data (germ cell versus somatic cells) and iii) the definition of a recombination hotspot (DSB versus recombination events) may explain this discrepancy.

Our methodology produces reliable RT profiles (supplementary information) and paves the way for similar experiments in which RT can be determined for other samples with limited number of cells, in particular *in vivo* cell populations. As far as we know, RT profiling of *in vivo* vertebrate cells was done only in zebrafish (Siefert et al. 2017); this is the first time it has been performed in mammalian cells. This technique is especially relevant in the field of cancer, in which it was shown that using the correct tissue of origin RT can best explain mutation rate (Polak et al. 2015; Supek and Lehner 2015). Currently, there are no RT profiles of primary tumors and the association between RT and mutation rates has been based so far on tissue culture cells. Measuring RT from *in vivo* tumors may help elucidate the correct mutation rate and aid in understanding the mutational spectrum in a given cancer.

In summary, by optimizing the RT profiling methodology we were able to determine the RT of two types of mouse germ cells. These novel RT profiles allow the identification of the tissue of origin of many genomic features. Moreover, they suggest a fundamental role for RT in determining multiple facets of genomic composition. Further research is needed for understanding the precise mechanisms by which this is achieved.

## Methods

### Tissue culture

Mouse embryonic fibroblasts (MEFs) were cultured in DMEM medium (BI) supplemented with penicillin, streptomycin, L-Glutamine and 20% v/v heat-inactivated (56°C, 30 min) FBS (BI). L1210 were cultured in L-15 medium (BI) supplemented with penicillin, streptomycin, L-Glutamine and 10% v/v heat-inactivated FBS (BI). Cells isolated from the bone marrow of female *C57BL6* mouse (10 weeks old) were grown in RPMI 1640 media (Gibco) supplemented with 10% fetal bovine serum (Hyclone), penicillin– streptomycin (Gibco), L-glutamine (Gibco) and 50μM of β-Mercaptoethanol (Gibco) on irradiated ST2 feeder cells. IL-7 conditioned medium (collected from J558L-IL7 secreting cells provided by A. Rolink) was added to the cells to select for pre-B cell populations for 14 days.

### Isolation of PGCs

Oct4-GFP+/+, Sox2-GFP+/- and M2rtTA+/+ mice were obtained from Jackson Labs. Oct4-GFP+/+ mice and Sox2-GFP+/- mice were bred to M2rtTA mice and females were sacrificed on day 13.5 of pregnancy. GFP positive cells were isolated from E13.5 gonads of either Oct4-GFP+/+ or Sox2-GFP+/- mouse embryos, resulting a pure population of PGCs (Yabuta et al. 2006), according to the following procedure. Embryos were dissected in PBS under the microscope. GFP-positive gonads were chosen by observation in fluorescent microscope and 4-8 embryos were processed per experiment. Gonads and mesonephros were first dissected and then separated into single cells using trypsin (BI) and 700µg/ml DNAse (sigma), followed by neutralization using FBS (BI). Cells were washed with PBS (BI), and filtered through 35µm mesh into 5ml polystyrene tubes (BD). GFP+ cells were isolated using FACSARIA III (BD) using cold conditions, into new 5ml polystyrene tubes, and fixated as described below.

### Preparation and growing of SSCs

SSC culture was prepared from the testis of 4-7 days old F1 C57/Bl6 crossed with DBA male mice, according to Kubota et al. (Kubota and Brinster 2008) with minor modifications. Testis cells suspensions were obtained using trypsin (BI) and DNAse (Sigma). Thy1+ cells were isolated using magnetic microbeads conjugated with anti-Thy-1 antibody (Miltenyi Biotec). Cells were examined for their replenishment potential in busulfan treated NODSCID mice.

SSCs were seeded on irradiated MEFs and grown in StemPro-34 medium (Invitrogen) as described by Kanatsu-Shinohara et al. (Kanatsu-Shinohara and Shinohara 2010). Cells were supplemented with 1% FBS (BI), human GDNF 20ng/ml (R&D systems), human LIF 50ng/ml (PeproTech), human FGF basic 1 ng/ml (PeproTech) and mouse EGF 20ng/ml (PeproTech). Cells were cultured for 4-6 weeks and split every 5 days.

### Fixation

MEFs and SSCs were washed with ice-cold PBS (BI), detached using trypsin (BI), and neutralized using the growth medium. Cells were moved to a 5ml Polystyrene tube (BD). All following reactions until filtration were done in the same tube and samples were kept at 4°C along the entire process. Cells were gently washed twice with ice-cold PBS, and resuspended in 250µl cold PBS. PGCs were diluted with up to 250µl cold PBS. For all cells, 800µl 100% high purity EtOH (Gadot) was added dropwise while slowly vortexing. Cells were kept for 1h to 24h at 4°C.

### Propidium iodide staining

Fixed cells were washed twice with 1ml cold PBS and spun down at 500g for 10min at 4°C after each wash. Cells were resuspended in 0.2ml PI-mix (PBS with 50µg/ml Propidium Iodide (PI) (sigma) and 50µg/ml RNAse-A (sigma)) and filtered through a 35µm mesh into a new 5ml polystyrene tube (BD). In order to enhance cell recovery another 0.2ml PI-mix was added and filtered to the new tube. For higher amounts of cells, we kept a concentration of 2.0×10^6^ cells per ml of PI-mix. PI-stained cells were incubated for 15-30 min at the dark before sorting.

### Flow cytometry

Cells were sorted using FACSARIA III (BD) based on their PI- intensity to G1 and S phases (Yehuda et al. 2017), using a flow rate of 1. Sorted cells were collected into 1.5ml Protein-LoBind tubes (Eppendorf) and moved to ice.

### DNA elution

DNA was extracted using DNeasy-kit (QIAGEN) and eluted twice with 2×200µl of the kit elution buffer (AE). DNA was moved to a 1.7ml MaxyClear tube (Axygen) which is compatible with the PureProteom Magnetic Stand (Milipore). X1.8 Agencourt AMPure XP beads (Beckman Coulter) were used to lower the elution buffer volume and gDNA was eluted from beads in 50µl EB (Qiagen). DNA amounts were measured using Qubit dsDNA HS Assay Kit (Thermo Fisher Scientific).

### Sonication

Samples of 50µl gDNA were transferred to a microTUBE Screw-Cap (520096, Covaris). Sonication was performed in the M220 Focused-ultrasonicator (Covaris) using 50W, 20% Duty Factor at 20°C for 120s, in order to reach an average target peak size of 250 bp. Sonication was verified using the D1000 or D1000 High Sensitivity ScreenTape using the Electrophoresis 2200 TapeStation system (Agilent).

### Library preparation for whole genome sequencing

Library preparation was done similar to Blecher-Gonen et al. (Blecher-Gonen et al. 2013) with some changes. Briefly, Sonicated DNA was subjected to a 50µl end repair reaction using 1µl End repair mix (E6050L, NEB), cleaned by 1.8X AmpureXP beads, followed by a 50µl A-tail reaction using 2µl Klenow fragment exo- (M0212L, NEB). The products were cleaned by 1.8X beads and were ligated by 2µl quick ligase (M2200, NEB) to 0.75µM illumina compatible forked indexed adapters. Ligation products were size selected by 0.7X PEG (considering the PEG in the ligation buffer) in order to remove free adaptors. 12-19 cycles of amplification were performed by PFU Ultra II Fusion DNA polymerase (600670, Agilent) with the following Primers: P7 5’AATGATACGGCGACCACCGAGATCTACACTCTTTCCCTACACGAC 3’, P5 5’ CAAGCAGAAGACGGCATACGAGAT 3’.

Amplified DNA was size selected for 300-700bp fragments by taking the supernatant after using 0.5X beads (which removed fragments greater than 700bp) followed by a 1.0X beads cleaning (which removed remaining primers and adapter dimers). The final quality of the library was assessed by Qubit and TapeStation. Libraries were pooled and sequenced on NextSeq (illumina) for 75 bp paired-end sequencing, generating 10M reads per each library (60M per experiment for triplicates of G1 and S).

### Generation of RT maps

RT measurements were performed as described (Yehuda et al. 2017). Briefly, sequencing reads were mapped using Bowtie 2 software. Discordant reads and PCR duplicated were removed. Every 200 G1 phase reads were binned in order to establish genomic windows. Corresponding S phase reads were counted in order to determine an S/G1 ratio for each window. Ratio data was normalized by subtracting the mean and dividing by the standard deviation. Continuous data was smoothed and interpolated using the Matlab csaps function (10e-16) at a resolution of 100kb (approximate average size of the windows). Continuous segments containing under 15 informative windows were removed from the analysis.

Published Replication Timing profiles were obtained from replicationdomain.com accessions: Int26004257, Int62905691, Int3190888, Int20705995, Int93235019, Int83562596, Int52548116, Int87752970, Int17857752, Int62150809, Int88652090. Data was smoothed and interpolated similar to smoothing of RT profiles generated in the lab.

### Determination of Differential Regions

Differential regions were determined using the likelihood ratio test at each window. The null model assumes that at a given point, all six measures come from the same distribution with a given mean. The alternative model assumes that the replicates of each sample belong to two separate distributions each with its own mean. Probabilities were calculated using the normal distribution probability density function. The variance used was estimated as the average of the normally distributed genome-wide variance of each sample.

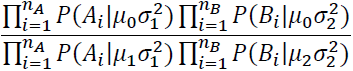

A chi squared p value was calculated for E2ln value of the ratio with 1 df. Bonferroni correction was used to control for multiple testing. All regions with a corrected p value below 0.01 were selected as differential and extended until the corrected p value exceeded 0.05.

### Statistical analyses

In analyses including multiple datasets, RT data was filtered to only include windows containing informative data in all datasets. In addition, sex chromosomes were excluded from the analyses, resulting in approximately 20,000 windows or 2Gb. Where applicable, RT data was filtered to include only the RT switching regions as determined by Hiratani et al. (Hiratani et al. 2010). Clustering was performed using the python seaborn clustermap using the correlation metric and the average method. Correlations were calculated according to Spearman and confidence intervals were calculated by bootstrapping the data (n=1000). LINE, SINE and GENEID data were obtained from the UCSC genome browser. Gene content was calculated as the percentage of bases covered by genes (from start to end of transcription) for each window. For mutation data we used mouse-rat diversity data which exons, splicing junctions and CpGs were excluded as described (Chen et al. 2010). Recombination hot spots are taken from (Lange et al. 2016). All genomic features were analyzed in 100Kb windows besides for recombination hot spots which are sparser and thus were analyzed in 1Mb windows. Chromatin accessibility was downloaded from the GEO database (GSM2442671, GSM2098124, GSM1014153, GSM1014149) (Yue et al. 2014; Guo et al. 2017; Li et al. 2017) and calculated in 100kb windows.

PCA was performed using the python sklearn PCA function. For experiments utilizing fixed GC content or chromatin accessibility, the relevant feature was sorted and split into four equally sized groups for further analysis. RT bins containing fewer than 50 or 15 windows for the 100kb windows data or megabase windows data respectively, were removed from analysis. Multiple regression analysis was performed using the python statsmodels OLS function. Autocorrelations were performed using the plot_acf function from the python statsmodels package. Partial correlations were calculated using a custom script based on the Matlab partialcorr function.

## Data Access

The data have been deposited in NCBI’s Gene Expression Omnibus (Edgar et al. 2002) and are accessible through GEO Series accession number GSE109804.

## Acknowledgments

We thank Dr. Dan Lehmann and Dr. Eleonora Medvedev for assistance with the FACS. We thank Dr. Idit Shiff, Dr. Abed Nasereddin and Alexia Azoulay for help in sequencing. We thank Prof. Yuval Dor, Dr. Tommy Kaplan and Prof. Eran Meshorer for fruitful discussions. We thank Dr. Micha Ben-Zimra and the Amos Tanay lab members for assistance in library preparation improvement and consultations. We thank Prof. George Iliakis for the MEF cells. We thank Dr. Shai Carmi for assistance in establishing the likelihood ratio test. This research was supported by the Israel Science Foundation (grant no. 184/16 to IS), the ISF-NSFC joint research program (grant No. 2555/16), the ERC Starting Grants (#281306) and the Israel Cancer Research Fund (ICRF).

